# Vibrational signatures and mimicry in ant-termite and termite-termite interactions

**DOI:** 10.64898/2026.03.15.711937

**Authors:** Sebastian Oberst, Joseph C.S. Lai, Theo Evans

**Affiliations:** Centre for Audio, Acoustics and Vibration, Faculty of Engineering and IT, University of Technology Sydney, Broadway, Sydney 2007, NSW, Australia; School of Engineering and Technology, University of New South Wales, Northcott Drive, Canberra 2000, ACT, Australia; School of Animal Biology, The University of Western Australia, Perth, Western Australia 6009, Australia

## Abstract

Eusocial insects fascinate researchers with their sophisticated communication systems and sensory specialisations. Ants and termites have coexisted in a long-standing predator–prey arms race, offering insight into the interplay between ecology and evolution. The subterranean termite *Coptotermes acinaciformis* can detect the predatory ant *Iridomyrmex purpureus* through footstep-induced vibrations, triggering defensive responses. Ants produce noisier walking signatures than termites, while the inquiline termite *Macrognathotermes sunteri* walks more quietly than its host, suggesting species-specific vibroacoustic strategies. Using statistical analysis of video-tracked motion and footstep vibrations in confined arenas across six ant and ten termite species, we show that *C. acinaciformis*, despite its body size, moves more smoothly than ants, which alternate between directed and erratic paths. Inquiline termites, by contrast, displayed erratic movements. Ants consistently produced stronger vibrations closely linked to body mass, while Highly Comparative Time Series Analysis revealed termite motions approaching chaotic dynamics. Notably, while *C. acinaciformis* and *I. purpureus* produced distinct vibrational signatures, *M. sunteri*’s signals overlapped with its host, consistent with vibroacoustic mimicry. Although the ecological nature of this association remains unresolved, our findings underscore the central role of vibrational cues in shaping interspecific dynamics and highlight vibroacoustic communication as an underappreciated driver of social insect ecology and evolution.

## I. INTRODUCTION

Eusocial insects such as Hymenopterans (ants, wasps, bees) or Isopterans (termites) interest researchers concerned with collective behaviour and communication [1–3], multiple object tracking [4], self-organisation of complex systems [5], insect vision and the theory of optical flow[6–8] as well as the design of insect-inspired hexapods [9–11].

For eusocial insects, communication among nestmates is an important aspect of information gathering. Vibrational information is perceived from substrate responses and may be intercepted by, e.g., parasitic butterflies, competitors, and predators [12–15]. Vibrations produced through feeding, e.g., on plant matter, tapping, or caused by motion such as simple walking are common [16]. Termites belong to the clade within cockroaches (Blattodea) and are among the most astonishing eusocial insects since they are blind and deaf to airborne sounds but listen to minuscule substrate-borne vibrations to make foraging decisions, to eavesdrop on competitors, and to drum alarm [14, 17, 18]. One of the most prevalent, yet possibly among the most difficult to analyse cues is that of substrate walking responses due to the smallest amplitude and large variability [15, 19]. Vibrations due to insect movement have been used to quantify the activity levels of the ant *Iridomyrmex purpureus* responding to different stimuli (illumination, stones, and neighbouring nestmates) [20]. Oberst et al. [15] showed that *Coptotermes acinaciformis*, a dominant subterranean termite species, can detect predatory ants (meat ants *Iridomyrmex purpureus*) based on their footstep vibrations only and that termites are significantly quieter than their predators, with the termite alarm pattern resembling the sequence, amplitude, and frequency of ant footstep responses, indicative of evolutionary adaptation and acoustic mimicry in termites.

Termites and ants have been in a predator-prey relationship for millions of years [15, 21], but their walking characteristics, including differences and similarities, are yet to be explored. Further, Oberst et al. [15] hypothesised that the soil-/litter-feeding termite *Mac. sunteri* is inquilinistic to *Co. acinaciformis*, which renders this a tri-trophic relationship as a predator-prey/host-inquiline scenario. This scenario is visualised in Fig. 1 which shows that while ants are omnipresent walking across a wood stump where termites are foraging, termites can just be found millimetres away from their predators [15]; the ants cannot perceive the termites, while the inquiline species likely can detect both the host and the inquiline based on vibrations only.

**FIG. 1.**
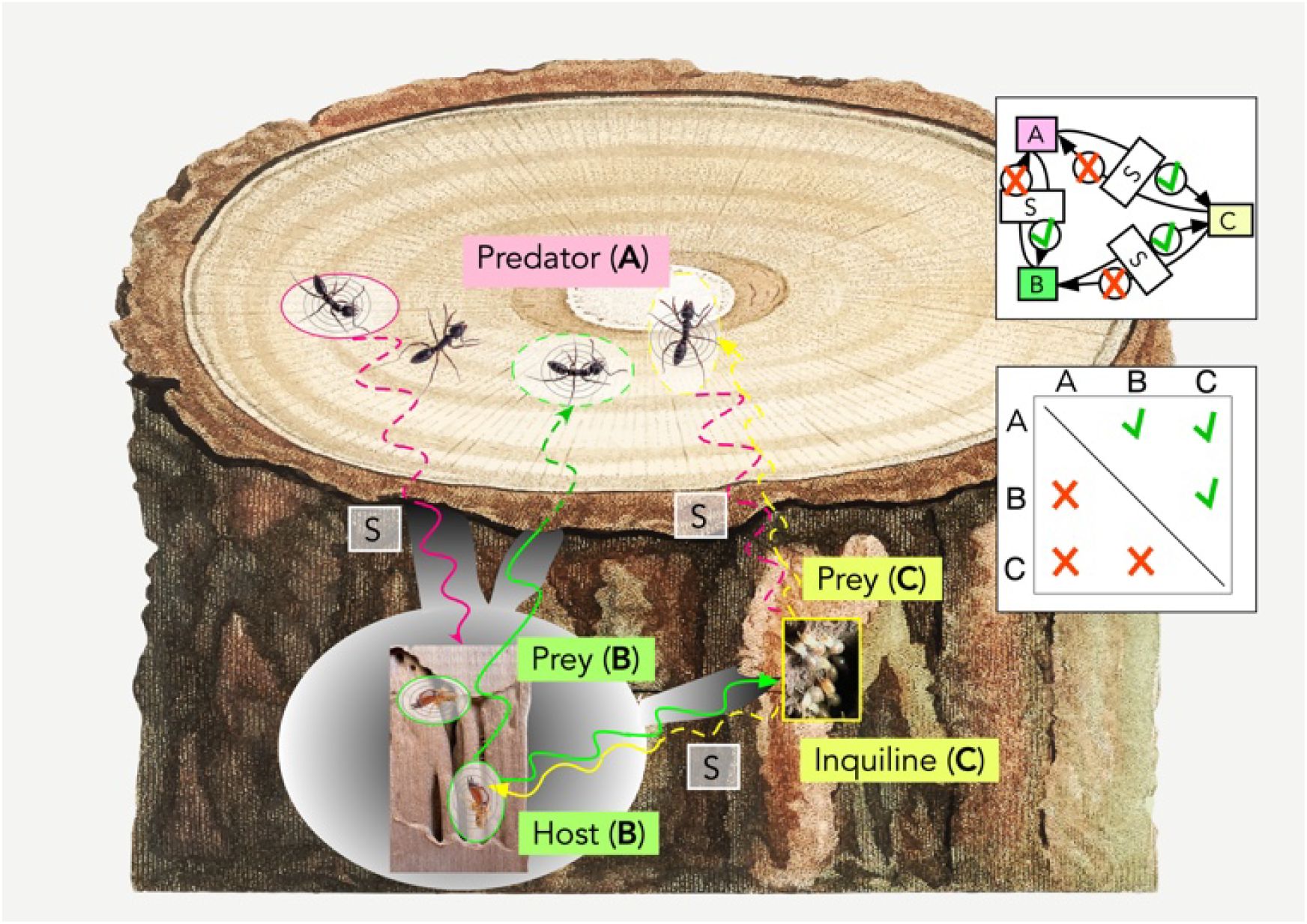
Conceptual diagram displaying the hypothesis based on [15] of an ant-termite predator-prey-inquiline relationship. Ants (A), predators, cause substrate-borne vibrations which termites(B), prey, perceive assisting them to dodge the predator; the prey termites cause acoustic emissions through the chewing of the substrate wood (S) and motion within their tunnels, which ants do not seem to hear [15, 22]. At the same time, the inquiline (in the host’s nest) eavesdrops on the host’s vibrations and those of the predator, and moves stealthily within the host nest without being detected. Insets show the communication pattern and a matrix (X is perceived by Y) indicating that the inquiline can perceive all vibrations, but A cannot hear the termites (photo credits: Sebastian Oberst; schematics freepik.dom).

Comparing the kinematics between ants and cockroaches, *Blaberus discoidalis*, ran faster with 25 body lengths per second (bl/s) [23] faster than ants, which showed for *Formica pratensis* 9 bl/s, or *Formica polyctena* 12 bl/s [24], yet only half as fast as *Periplaneta americana*. However, while their gait was found to be rhythmically fluctuating the same way, differences in movement control are apparent, presumably due to ants moving in more rugged habitats compared to cockroaches being runners on flat terrain [24]. Reinhard et al. [24] used a high-speed camera (992 frames/s) and a miniature force plate to analyse the motion of red wood ants and verified a typical tripod pattern (in-phase ground contact of the tripod). Reinhardt et al. [24] hence suggested that wood ants permanently cling to their substrate in a climbing style even on level surfaces (*level climbing*). Reinhardt and Blickhan [25] further studied wood ants and showed an unaltered tripod gait pattern over a wide range of speeds (controlled via stride length and frequency). The ants’ front legs showed high compliance, which reduced vertical oscillations and increased stability by maintaining ground contact, enabling the possibility of bouncing gaits without aerial phases to perform grounded running.

The ground reaction force and gait cycles have been studied early on, assuming running/hopping or walking gait, reducing the tripod action to bipedal and monopedal motion in the sagital plane (based on a spring-loaded inverted pendulum (SLIP) model) [26] or a lateral-leg spring (LLS) model in the coronal/horizontal plane [27]. Oberst et al. [19] developed a signal processing method to deconvolve inversely and blindly the ground reaction forces as the underlying excitation of the substrate response based on only measuring substrate vibrations. They found that the vibration response also contained different step functions related to the variation in impact time of the individual legs.

While the walking of ants has been classified following a Brownian motion or random walk (Lévy) process [28, 29], not many studies are concerned with a more detailed statistical analysis comparing different species. As indicated by Pyke it is time to understand the movements of organisms and therefore to abandon the Lévy foraging hypothesis [30], but there is no comprehensive comparison between ant and termite walking across different species and their resulting footstep vibrations. Neves et al. [31] studied the video-tracked trajectories of the ant *Pheidole rudigenis* (major and minor workers) and the mealworm beetle (*Tenebrio molitor*) using recurrence plots and found deterministic, quasiperiodic traits in ant patterns. Similar to recurrence plots and recurrence plot quantification measures [32, 33], the highly comparative time series analysis has been developed as a massive feature extraction tool for effective clustering of complex dynamical data [34].

Despite extensive literature on ant and cockroach locomotion and gait mechanics [23–27], surprisingly little is known about how walking kinematics translate into substrate-borne vibrational signatures across species, and how these signatures differ systematically between ants, host termites, and inquilines. Walking-induced vibrations are challenging to analyse due to their low amplitude, high variability, and sensitivity to substrate properties [15, 17], yet they represent one of the most pervasive signals available for eavesdropping in subterranean and plant/wood-based habitats [35, 36].

Here, we investigate whether walking-induced vibrational signals and associated motion patterns differ consistently between ants and termites, and whether these differences align with ecological roles such as predation, host status, or inquilinism. We combine high-resolution video tracking of locomotion with laser Doppler vibrometry to quantify motion characteristics and substrate vibration responses in six ant and ten termite species. To compare complex time series of walking-induced vibrational signals across species, we employ highly comparative time series analysis (HCTSA) as a feature-extraction and clustering framework, enabling a systematic assessment of similarities and differences in signal structure.

Our specific aims are threefold: (i) to compare walking kinematics and spatial movement patterns between ants and termites across multiple species; (ii) to quantify differences in the magnitude, temporal structure, and spectral properties of walking-induced vibrations; and (iii) to assess whether the vibrational signatures of the inquiline *M. sunteri* overlap with those of its host *C. acinaciformis*, consistent with vibroacoustic similarity or convergence. Rather than testing adaptive mimicry directly, we examine whether the observed signal patterns are consistent with ecological hypotheses of detectability, stealth, and coexistence in ant–termite systems.

## II. MATERIALS AND METHODS

### A. Collection of species

For this study, six ant and ten termite species were collected. For ants *Iridomyrmex purpureus, Lasius niger, Papyrius nitidus, Camponotus aenopilosus, Camponotus consombrinus* and *Rhytidoponera metallica* were collected from three colonies at sites near the Mt Pleasant, ADFA, Canberra (Cb) (35°29’18” S, 149°16’61” E, ACT, Australia) and from sites in the Binya State Forest, Griffth (Gr) (34°21’04” S, 146°25’94” E, NSW, Australia).

For *I. purpureus* a wild form and a laboratory form were used, indicated through * and ** in the following. Three laboratory colonies were reared one year before the experiment started, with up to 187, 201, and 240 individuals. The ants reared in the laboratory were young colonies and expected to have smaller built individuals due to colony maturity, non-ideal rearing conditions, and constant but elevated temperatures [37, 38], details see Supplementary Materials. Having smaller ants of the same species available would serve as a natural benchmark model to determine, in principle, whether their extracted features were identical to the wild form.

For termites *Mastotermes darwiniensis* (12°27’57”S 131°03’09”E, Darwin (Dw), NT, Australia), *Cryptotermes secundus* (*Coptotermes acinaciformes* (Southern form (Gr) and Northern form (Dw)), *Co. frenchi* (Gr), *Schedorhinotermes reticulus* (Gr), *Macrognathotermes sunteri* (Dw) *Microceratermes nervosus* (Dw), *Tumulitermes pastinator* (Dw) and *Nasutitermes exitiosus* (Gr) were collected from three colonies each. We did not rear a laboratory form.

### B. Setting up the experiments

#### 1. The test rig

The experiments were conducted in an anechoic room (150, Hz cut-off frequency, dimensions of 3.5 × 3.5 × 3.5, m^3^) with a controlled temperature of approximately 30°C maintained using an oil fin heater (SUNAIR - Sunair Oil Heater, 11 Fin Fan, thermostat, 2400, W). Humidity was left uncontrolled but remained relatively constant at a low level of 28 ± 5% relative humidity (RH). This low RH ensured minimal moisture absorption, thereby reducing damping in the wood [17, 39]. Temperature and humidity measurements, along with video recordings, were synchronized using an Arduino UNO microcontroller equipped with a temperature and humidity sensor (Freetronic DHT22) and a switch connected to an LED indicator, which provided visual recording status in the videos. To prevent overheating, the laser vibrometer was actively cooled with Peltier elements (European Thermodynamics Peltier Module, 17.2, W, 4.59, A, 7.5, V, 62 × 62, mm^2^). These elements were mounted beneath a frame and connected to cooling fins and a steel plate attached to the underside of the vibrometer using thermal grease.

The experimental setup (lid only of the insect box used in Oberst et al. [15], foam, wood on two bricks), schematically illustrated in Fig. 1, was supported by foam on a steel frame positioned at the centre of the anechoic room. Residual environmental vibrations transmitted through the flooring and omega springs (resonance frequency of 8, Hz) of the anechoic room were detectable and attributed to personnel walking past or the impact of closing steel doors in adjacent workshops. To isolate the steel frame and setup from these vibrations, an air-cushioned passive vibration bench top (Kinetic Systems, Boston, MELpF, 24, kg) was employed. The bench was calibrated at 2, MPa and placed atop a vibration mat on concrete slabs weighing 80, kg. Calibration involved measuring the impulse response (PSD and amplitude) from a 5, kg weight repeatedly dropped from 0.5, m height onto the concrete floor outside the anechoic room, at 10 points forming a semicircle approximately 1.5, m from the room’s outer shell. The location with the highest response sensitivity was used to adjust the pressure of the air-cushioned bench top, minimizing external impacts so that the transmitted power per frequency from dropped weights was at most the same order of magnitude as the noise floor.

The lid of the insect box consisted of a rectangular PPE container lid glued to a cylindrical container with a radius of 25, mm. A veneer disc was placed inside the cylindrical container and secured with a PVC tube (cf. [15, 20] for more details). Veneer discs were sourced from *Pinus radiata* veneer sheets intended for plywood panel production. The specific veneer disc used in this study had the following material properties (cf. Set B [40]): an area of 2, 845, mm^2^, a weight of 1.3564, g, moisture absorption of 12.87% relative to dry weight when exposed to 80% RH, a mode skewness of −0.390 (measured via light intensity distribution in digital images), and an average thickness of 0.94, mm (measured 1, cm from the edge every 120°) [17]. The wood grain was oriented in-plane with the veneer disc, and its direction was consistently aligned with other components (box, container lid, spacer) of the insect box. To prevent insects from climbing the PVC tube and inducing unnatural behaviour, two thin layers of Fluon (a polytetrafluoroethylene-based coating) were applied to the tube walls, as described in Oberst et al. [20].

#### 2. Handling of insects

After collection, the insects were stored in an environmental room with constant temperature and humidity (about 28°C, 80 ± 5 % RH) and then manipulated for the experiments using feather-light forceps with rounded tips (Ento-supplies, E122BT-10). Ants were cooled down to about 10°C for ca 2 min. and weighed using an AEG scale with four-digit accuracy. The insects were placed on the thin veneer disc setup using the setup depicted in Figure 2 (a) in the anechoic room (about 30°C, 28 ± 5 % RH), then left for about 20 min in the anechoic room to acclimate for temperature and humidity differences.

**FIG. 2.**
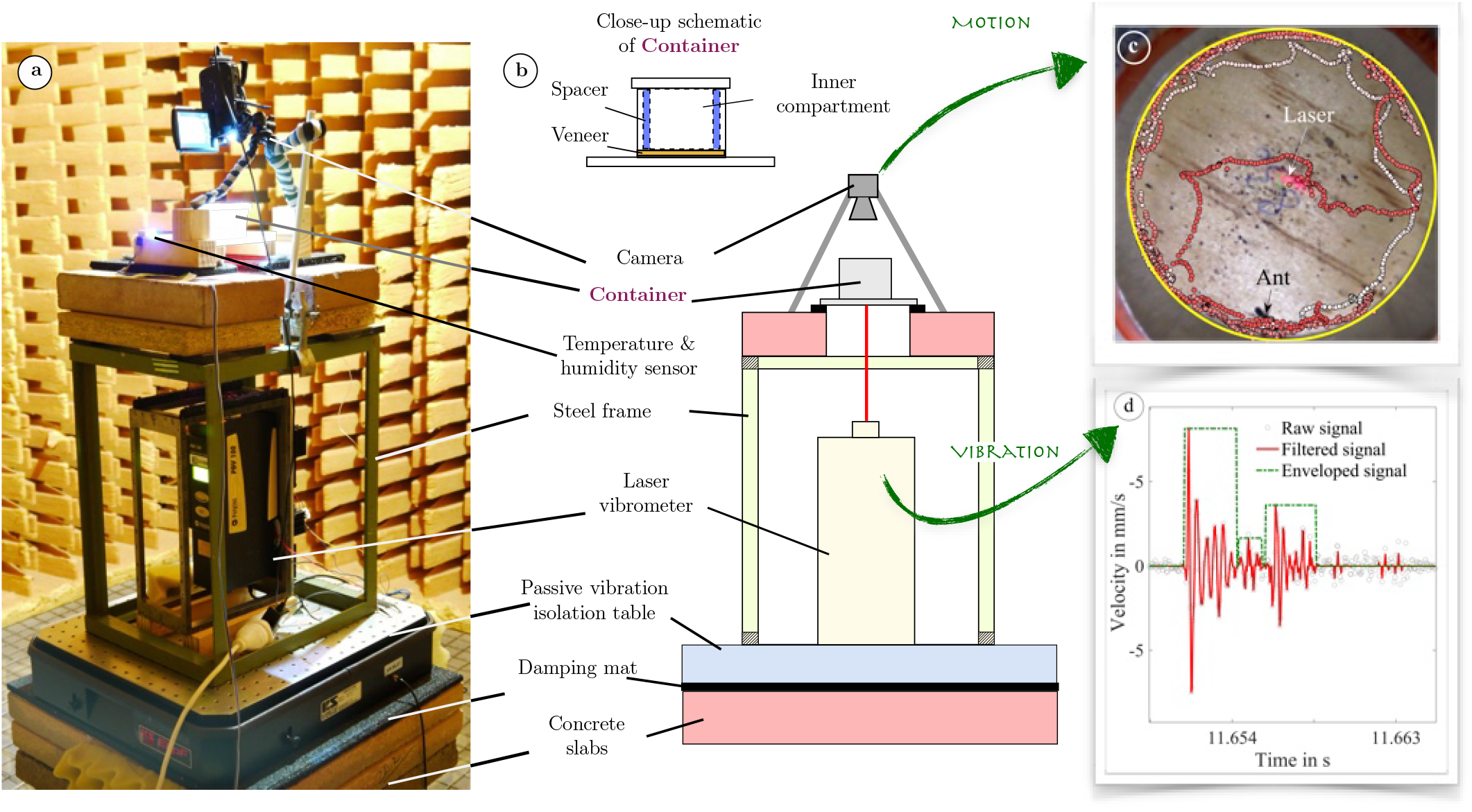
Experimental setup and schematic to monitor motion and vibration of ants and termites. (a) Experimental setup in the anechoic room and (b) schematic with a close-up of the cylindrical container. (c) The results of the tracking algorithm for one ant *Iridomyrmex purpureus*, and (d) Raw, filtered, and enveloped signal of an individual ant of *I. purpureus* walking on the veneer disc. (photo credits: Sebastian Oberst)

**FIG. 3.**
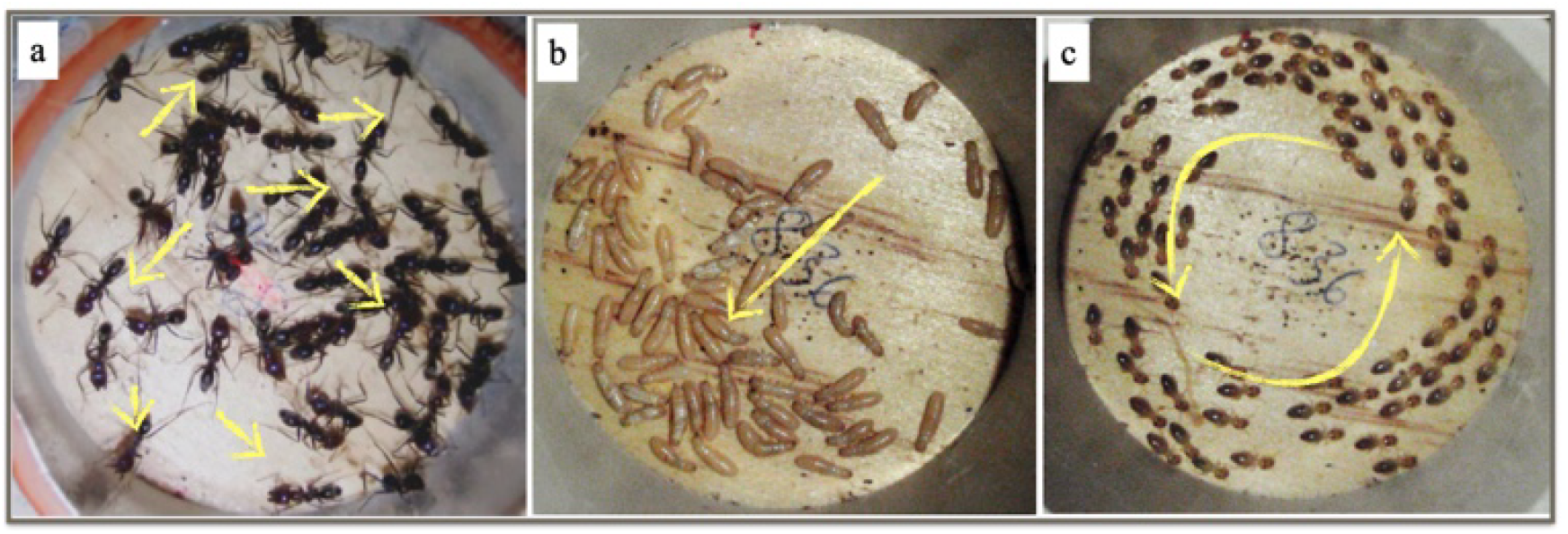
Preliminary observations made in a container. (a) *I. purpureus*, (b) *Cr. secundus*, and (c) *N. exitiosus* walking in the inner compartment of the insect box. It was observed that the ants’ walking behaviour is more erratic than that of termites, which follow each other either (b) in 2-tandems or (c) n-tandems in closed circles.

### C. Motion analysis

While the insect vibration was recorded using the LDV, the individual insect’s motion in the arena was monitored using a high definition (HD) digital video camera (BENQ M23 Full HD), attached to a 1/4” aluminium bar, mounted on a flexible mini tripod bracket (mini octopus tripod, Gadgets 4 Geeks). Care was taken to record directly above the container at a distance of 120 mm, far enough to have acceptable image distortion. However, due to the close-up recording, the circular walking area became slightly elliptical. The videos were down-sampled to 377 × 141 pixels and saved as *.png files, then analysed using an ant-tracking algorithm based on Kalman filtering [41].

#### 1. Preparation of the video files

To analyse individual ants walking, the videos were segmented into recordings of one minute length triggered by the LED[42]. The video files were read in using Matlab 2020A. Each video file was calibrated manually by selecting (1) the starting point (centre of veneer), which will have the elliptical ant arena inscribed, and (2) two position vectors to control the direction of the wood fibre. Then, an elliptical arena (inverted elliptical mask) was superimposed onto the images, which were rotated to obtain the same direction of the fibre and saved as a monochrome 8-bit PNG file relative to the number of the veneer disc and the grain direction. The mask had two purposes: (1) to reduce the file size to only the area of interest and (2) to cut away parts of the PVC walls that interfered with the video tracking algorithm through reflections. Fluon, which was applied to the walls of the PVC spacer to prevent the insects from climbing up the walls, made the PVC surface, which could not be removed, slightly more reflective so that in some instances the ant-tracking algorithm would not only track the ant but also the ants’ mirror-images. This had minimal effect on the overall quality of the tracking in a statistical sense, but had to be manually removed when concluding individual traces.

#### 2. Tracking the insects

The ant-tracking algorithm employed uses the work of Wauthier [41] (which is based on Stauffner and Grimson [43]), and is developed to detect and track foreground objects; here specifically their code was adapted to reliably track the insect, including a moving position vector within an elliptical stencil (with major and minor axis inscribing the insect) and its statistical weighting. A mixed Gaussian background adaptation algorithm with Kalman tracking was used. The background adaptation algorithm is employed to distinguish between objects moving in the foreground and background pixels, while the Kalman tracking algorithm is employed to determine the trajectory of a moving object with the information of the foreground pixels [44]. Based on the Bayesian principle, a hypothesis (posterior) is updated using the model information and a prior probability. For foreground detection, a background model captures the distribution of previous pixel data (a mixture of isotropic Gaussians [45]). The distribution properties (mixing proportions, means, and variances) are updated, and a verified background model is compared to each pixel to determine foreground and background membership degree. Foreground objects are then detected by connecting all foreground pixels to blobs [41, 43]. For each of the foreground objects, the geometrical centre was determined by applying weighted averaging, and then the centre was stored in a polar coordinate matrix [46].

#### 3. Postprocessing

A result of a tracked ant of the species *I. purpureus* is depicted in Fig. 2 (c), the darker the colour, the more recent the recorded event occurred. The tracking algorithm detected the motion of a single ant using 30 frames per second (fps) and statistical weighting of the direction vector within an elliptical stencil. The step width, the velocity, and from it the maximum speed per second, the speed in body length, the activity, and the total distance traveled (based on average speed) are recorded. The length of an individual was estimated using the number of pixels, which was found to be well correlated with the insect’s weight (see Figure S1, Supplementary Materials). The tracking algorithm only failed to record activity if the insect was not moving (e.g., while grooming). The relative frequency of insect motion over the area was recorded to calculate the distributions (*n* = 4). To describe the overall motion regime of the insects qualitatively we decided on three classifications, (1) *rotational motion*, if more than 30% of the relative frequency of tracked motion was distributed within a certain (arbitrary) radius of the arena’s veneer disc and if less than 10% of the relative frequency were found in the centre disc (radius 10mm); (2) *mixed motion*, if either of the criteria in (1) was violated, and (3) *seemingly random* where both criteria were violated.

### D. Vibration measurements

The measurements were conducted using a single-point laser Doppler vibrometer (LDV) Polytec PVD-100 with a data acquisition system Polytec VIB-E-220, connected to a Toshiba notebook with Vibsoft 8.8 analysis software. The laser vibrometer only measured vibrations of the centre of the veneer disc. By inspecting video recordings, only segments of vibration signals measured with an individual ant walking within the ant’s length of the measurement point were analysed. The insects were observed for fragments of 172.8 s at a 12 kHz sampling rate. The motion of the specimen in the container (Fig. 1) was recorded using a digital video camera BENQ M23 (Full HD 1080p, 30 fps) to later correlate qualitatively and quantitatively the average activity level and the behaviour to detected vibrations for a larger study. The inside of the container was illuminated during recording using the camera’s built-in cool white LED (3 mm). Reference measurements with/without insects were taken before each measurement to determine the noise floor (average RMS value of about *σ* =104 *µ*m/s).

### E. Statistical analyses

Based on the motion analysis and the vibration recordings, a statistical analysis is conducted. For all analyses, unless otherwise indicated, only a single ant (*n*_*i*_ = 1) is studied, of which *n*_*s*_ = 4 samples are chosen.

#### 1. Distributions

##### a. Motion analysis

Analysis of the video recordings was performed for 60 seconds after discarding the initial 100 seconds. The distributions for the motion analysis were generated by spatially averaging the trajectory of individually walking insects for 1800 frames. The data was then normalised by the highest occurrence per species to estimate the relative frequency. The detailed walking parameters are quantified by the median speed (in body length (bl) per s), the median absolute speed (mm/s), and the average activity in per cent of frames where movement is detected relative to the overall frame number. The first 100 seconds of the recording were clipped, as was the remaining time after the 60-second analysis time.

##### b. Vibration analysis

As indicated in Fig. 2 (d), the data were recorded using a vibration-isolated laser vibrometer. The measurements were nonlinearly filtered using the *ghkss* geometric filter of Kantz and Schreiber [47] with a fixed embedding dimension of *m* = 3, a delay *d* of unity, a manifold *q* = 4, and *i* = 8 iterations. We found that the dimension for the signal at hand never exceeded 3, and that 1 as a delay was good enough, presumably due to the noise content of the unfiltered signal and sometimes longer interpulse intervals when the insect was standing still. The ghkss filter enables significant improvements in signal-to-noise ratio, eliminating almost completely the noise as shown in Fig. 2(d), cf. [2, 15, 17, 19]. The signals’ single-step responses are isolated using a signal enveloper, which determines the averages of amplitude and the decay times of ant footsteps by generating a rectified square wave. For this purpose the standard deviation *σ* of the background noise signal 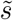 corrected by its offset is taken as the steady-state (*ss*) error as reference signal forming a tolerance and signal *ss* ± *σ* with 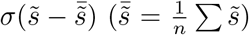. For 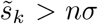, with 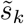 a square wave of magnitude *M*_*k*_ is generated. The length of the square wave *λ*_*k*_ (generally the exponential decay constant) is determined by counting the incidences *I* of zero-crossings within the settling time *t*_*s*_. For *I* ≡ 1, a zero-square wave is generated which is merged with all following *I* ∈ {0, 1}. Due to the different nature of signals and according to the ‘Empirical rule’ [48] the ant signal has been selected to exclude 99.7% (3*σ*) of the background noise signal; while the termite signal has been selected to exclude 95% (2*σ*) of the background noise.

The *M*_*k*_ and *λ*_*k*_ were sorted and normalised concerning their largest frequency of occurrence row-wise in a matrix, the median value of each distribution in each column (bin) was estimated, and a curve connecting the median points was fitted, cf. Figure S2 for details.

From the distribution of attenuation time and amplitude for ants and termites, a correlation analysis for their means is conducted after estimating the distribution properties and the median as a position parameter; the weight and size of ants are taken from Figure S1. As a position parameter, we used the median for the vibration and for the attenuation time 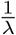, *λ* representing the rate parameter of the distribution, i.e., events per unit time of an exponential distribution.

#### 2. Power Spectral Density

We calculated the power spectral density (PSD) for each of the time series analysed. We applied an anti-aliasing filter with a Kaiser window to the time trace and used Welch’s method of averaging (Hann window with 128 samples) with cut-off frequencies of 150 Hz and 5 kHz [49]. The PSD, obtained by the Fourier transform of a time-varying function (in this case, the auto-covariance function of the walking vibration signal), expresses the energy per Hz in the frequency domain [49**?**]. The PSD is calculated by using

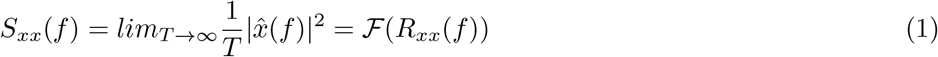

with ℱ being the Fourier transform operator and *R*_*xx*_ being the auto-covariance function (cf. Wiener-Khinchin Theorem [50]). With all PSDs computed, we then extract their statistical properties such as standard deviation and average peak power at their fundamental frequency. We also calculated the integrated power using a PSD integration analysis (PSDIA) similar to that in [51], but just using the ultimate integrated power and forming a statistic from it (*n* = 10), hence

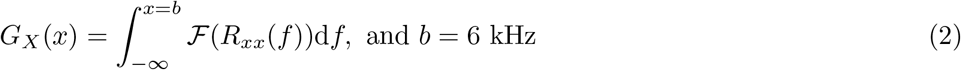

As a reference value, we use 200 *µ* m/s, ca. 2 times the standard deviation (2*σ* - covering ca 95% of the data points) of the background noise (rms = 104 *µ*m/s).

### F. Highly Comparative Time Series Analysis (HCTSA)

Highly comparative time series analysis (HCTSA) was introduced by Fulcher and Jones [34] to compare different time series toolboxes accumulating to extract up to 7,700 outputs (features)[52]. It allows comparing different time series and clustering them by extracting statistically significant main features of a time series [34]. We study the ability to cluster and therefore separate different time series, and we adopt a low-dimensional representation of the data using principal component analysis. In the HCTSA, the main matrix, the time series studied is the input in its very left column, followed by the actual output, the feature (HCTSA) matrix. The feature matrix contains about 7,700 features [34] and sorts them according to significance using a colour code. In a second graph, a reduced feature space is provided in the principal component analysis, and the different features related to time series are clustered with the first principal component (PC-1) and the second principal component (PC-2) in a scatter plot. PC-1 captures the maximum variance in the data. PC-2 captures the next highest variance, subject to being orthogonal (uncorrelated) to PC-1.

One can assume that if features are very different, clusters can be separated in the HCTSA matrix as well as in the scatter plot; if features are somewhat similar, they should be clustered as blocks in the HCTSA matrix but can show overlapping clusters in the scatter plot. If they are well mixed, no clear blocks or clusters are formed.

We quantify the group clustering ability by introducing a scoring system. The different data is then further studied for its ten most prominent features, which are then discussed.

#### a. Cluster scoring system

Using HCTSA, the clustering of features of a time series is not only studied by reducing the feature space to the two main (linearly independent) features, but also by the closeness (neighbourhood) of the full feature vector to related feature vectors in the HCTSA matrix. If a neighbouring time series belonged to the same kind of signal, it was counted as a cluster of size *c* = 2, if two time series were direct neighbours, it was counted a cluster of three and so forth, with *N* (the overall number of same species time series studied) being the largest cluster (cf. Oberst et al. [40]). The number of clusters received a weighting function *w* of its squared cluster size *w* = *c*^2^, which was then normalised by its potentially highest value which is 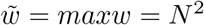 to generate a percentage score *Q*, representative of the quality of clustering (*Q* = 0 no clustering possible, *Q* = 1 perfect clustering (in the following *clustering-quality or C-score*).

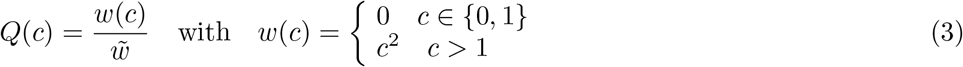

We then apply a model based on combinatorics and probability theory. The general goal is to determine if the relative clustering of direct and indirect neighbours [40] is a product of mere chance or not [53]. The combinatorial model applied is related to the question of “seating arrangements”, where groups of friends are seated together, either as individual blocks or multiple blocks, out of a larger cohort. The probability of having a certain configuration is derived from a relative frequency as an estimate formed from 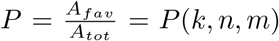, where *k* is the total number of species to be grouped, *n* is the total numbers of measurement, and *m* is the number of groups to be formed. *A*_*tot*_ is the total number of possible seating arrangements for *n* students, and *A*_*fav*_ is the total number of arrangements according to the question. One would ask, for instance, how many arrangements exist where three measurements of the same species are clustered together among all the other groups of measurements.

Hence, let *g*_*i*_ be the number of measurements of each species, then

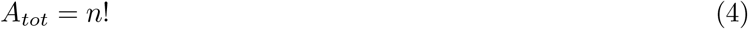

is the total number of ways to arrange *n* measurements in *n* locations, which will later be the denominator. Favorable (feature-based) arrangements depend on the specific groups to be identified. The general approach is to (1) partition the group of measurements into species. (2) Treat each group as a block that will be located as direct neighbours/sit together, and (3) arrange the blocks and other students within the arrangement.

1. The number of ways to partition *k* species into *m* groups with sizes *g*_1_, *g*_2_, …*g*_*m*_ is,

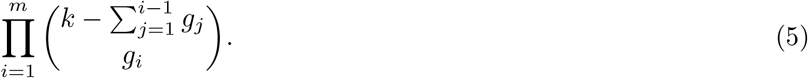
2. The number of ways to arrange the *m* groups is *m*! hence each group can internally be arranged in *g*_*i*_! ways and the total internal arrangements for all groups are:

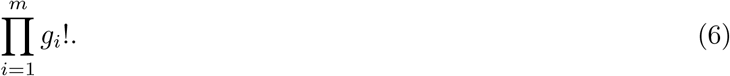
3. The number of ways to arrange the *m* groups and the *n − k* other students is (*n − k* + *m*)! so that the total number of “favourable arrangements” becomes

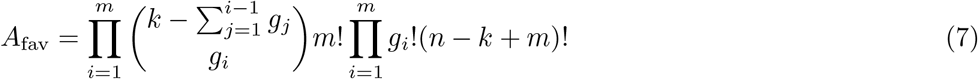

The probability of a given arrangement can therefore be estimated from the total number of favourable arrangements and the absolute total number of arrangements:

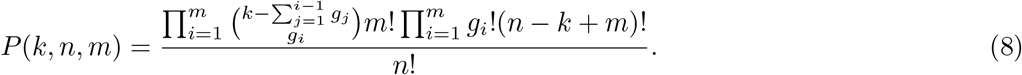

Both the clustering percentage according to *Q*(*c*) and the probability *P* (*k, n, m*) relative to the probability that the clustering is mere chance can be used as a metric of the quality of performance of the HCTSA in distinguishing walking vibration signals of insects.

##### 1. Time series used

Here, we first evaluate the functioning of the HCTSA tool by using benchmark functions and then applying it to measured data. We always generate a time series of 2,000,000 samples from which we extract a small time series segment of 12,000 samples (*N* = 5 each), corresponding to 1 s measurement time of experimental data. For live species, three colonies are compared, and only individual ants and termites are compared.

###### Case A: General benchmarks

The benchmark case compares computer-generated pseudo-random numbers (uniformly distributed white noise) against a background noise signal of the setup described in Figure 2 as candidates of high-dimensional deterministic or even random behaviour. The third signal is that of a Lorenz system in the chaotic regime, as a representative of a deterministic, low-dimensional system with a maximum prediction uncertainty, followed by a Lorenz system in a limit cycle regime (nonlinear system) and a sinusoid of 100 Hz and unit amplitude (simple periodicity).

###### Case B: Filtering

We then tested how the HCTSA algorithm works on filtered data compared to background noise and unfiltered (original) measurement data of an ant of the species *Iridomyrmex purpureus*. This is to examine how the quality of the signal influences the prediction and clustering quality. As a filter, we used the well-validated locally projective geometric *ghkss* filter algorithm of Kantz and Schreiber [47]), cf. with [19, 54].

###### Case C: Ants vs termites

We tested how the HCTSA algorithm works on filtered time series data of ant (*Ir. purpureus* and termite signals (*Co. acinaciformis*) in comparison to the chaotic signal and the background noise.

###### Case D: Size effect

To test whether the nonlinear filtering works well enough for maximally similar signal patterns and to test whether it removes enough noise, signals of ants (*Iridomyrmex purpureus*) reared in the laboratory are compared against the wild form of the species. The laboratory form is only about 65% as long (as heavy) as individuals caught in the wild [15] and represents, therefore, a valid mechanistic scaled model with presumably the same features and gait pattern.

###### Case E: Predator-prey/host-inquiline

In the Northern Territories, *Coptotermes acinaciformis* builds mounds, adjacent to a (commonly Eucalyptus) tree. Its mound is often also home to the mutualistic species *Macrognathotermes sunteri*, which builds its nest on top of *C. acinaciformis* mounds (cf. [3] for an photograph). While *Iridomyrmex sanguineus* is the dominant meat ant species in the north, *Iridomyrmex purpureus* has been sighted near Darwin [55], justifying the study of the difference between these three species. In Oberst et al. [15], footstep vibration responses of ants and termites were compared, and highlighted an important but under-explored ecological linkage [21]. *Coptotermes acinaciformis* is a dominant subterranean and tree-nesting termite in Australia’s south, while *Iridomyrmex purpureus* (meat ant), a diurnal and abundant, carnivorous ant species of the family Formicidae, is one of its major predators [15, 56].

###### Case F: All species in one HCTSA analysis

We eventually use all species and test whether we can cluster them according to a PCA analysis or the developed scoring system. Using a probabilistic model, we show how significant the findings are and whether clusters found are deterministic or rather based on chance.

## III. RESULTS

### A. Motion analysis

The initial motion analysis qualifies the walking pattern of individual ants.

As indicated in Figure 4, ants show in 6 out of 7 cases (85%) mixed motion, with almost a random pattern and direction reversal. Only *Rhytidoponera metallica* shows a predominantly circular pattern similar to a path following pattern [57].

**FIG. 4.**
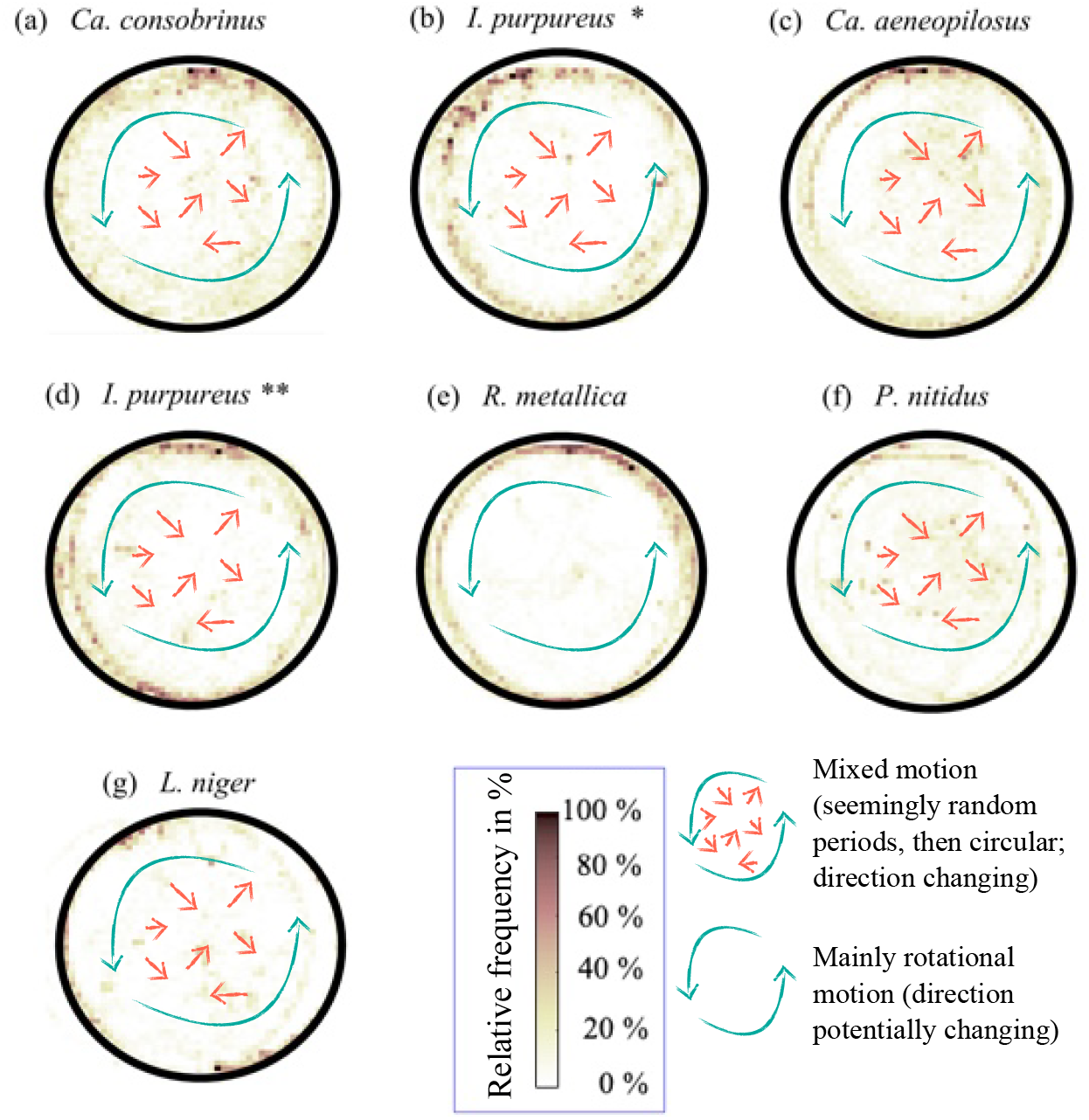
Ants’ spatial distribution - ants move in a seemingly random fashion. (a) to (g) Relative frequency occurrence of ants walking over the veneer disc analysed via video tracking algorithm [41, 44]. Six out of the seven recordings show a mixed motion, i.e., ants walking in circles along the inner edge of the PVC tubing insert and on the surface erratically. Only (e) *Rhytidoponera metallica* mainly moves about in a circular fashion (single ant only).

Figure 5 shows slightly different behaviour for ten cases of termites, with 40% showing mixed motion (a,d,g, and i), 30% (c, e, and f) having clear path following behaviour, and 30% (b, h, and j) showing a seemingly random behaviour. Dominant species, such as *N. exitiosus* or *Co. acinaciformis* tend to walk in a clear circular pattern. The seemingly random pattern is shown by *Mac. sunteri, Mic. nervosus* and *Cr. secundus*. The species, which seemed to walk the most randomly, is *Mac. sunteri*.

**FIG. 5.**
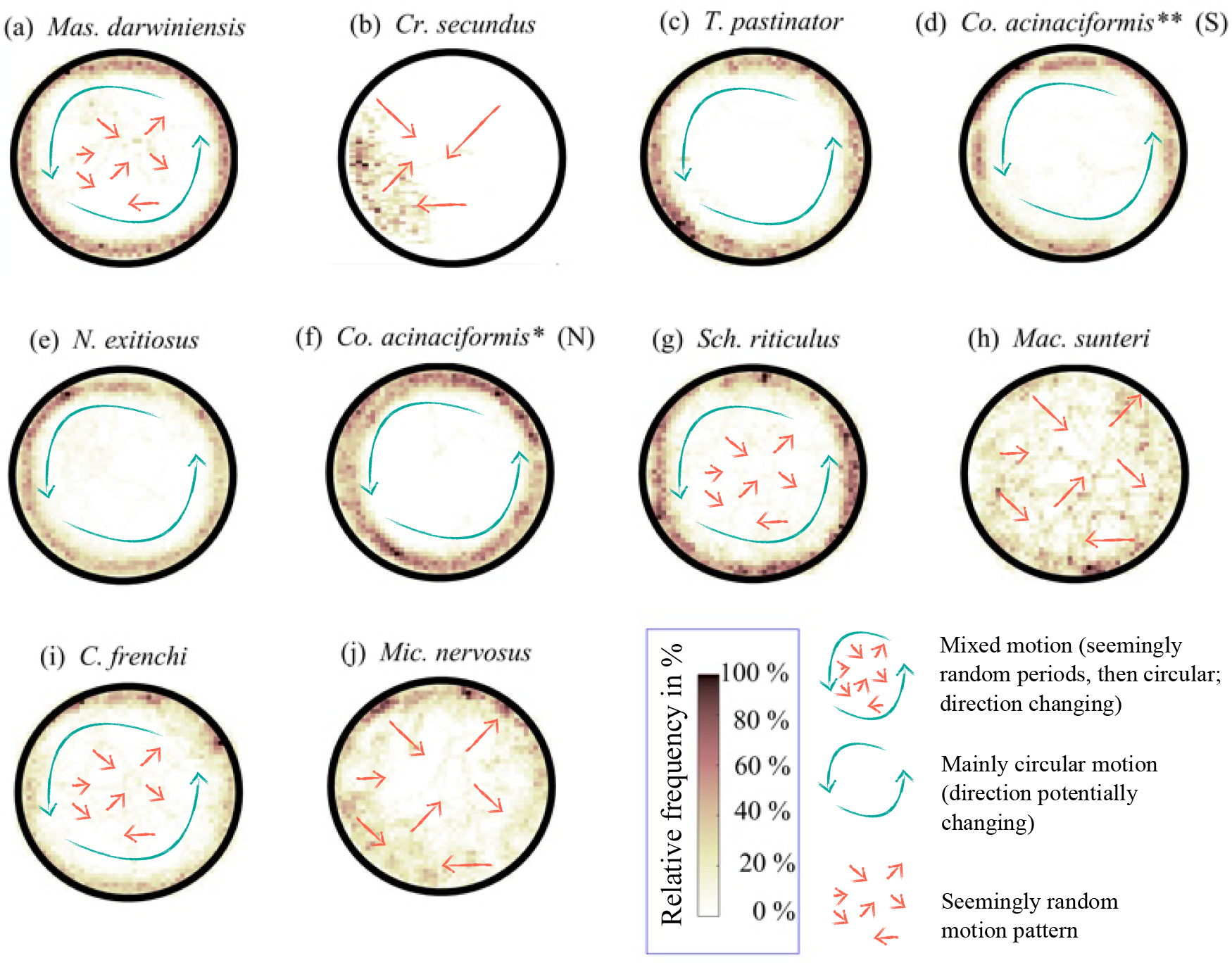
Termites’ spatial distribution - termites follow a pattern. (a) to (j) Relative frequency occurrence of termites walking over the veneer disc in the container as analysed via video tracking [41, 44]. Termites walking seems to be oriented and patterned; only four of the ten species studied (a, g, i) showed mixed motion (with stronger circular pattern than in ants); three species (c, e, f) followed very strongly a circular pattern and three species (b, h, j) showed highly irregular motion, which was the most irregular for *Mac. sunteri* and can be described as *clustered* irregular in case of *Cr. secundus*. (Note: subfigure (d) showed some artefacts at the edges where reflections of the wall caused more issues than in the other recordings due to slight misalignment of the camera.)

In Figure 6, the phase angle of motion for walking ants and termites was plotted as well (individual trajectory only). Differences previously observed are confirmed here as well - termites (including *Mac. sunteri, Cr. secundus* and *M. nervosus* walk more often and consistently in one direction and not change the path radius as often as ants, which change the path and do not show a clear direction reversal.

**FIG. 6.**
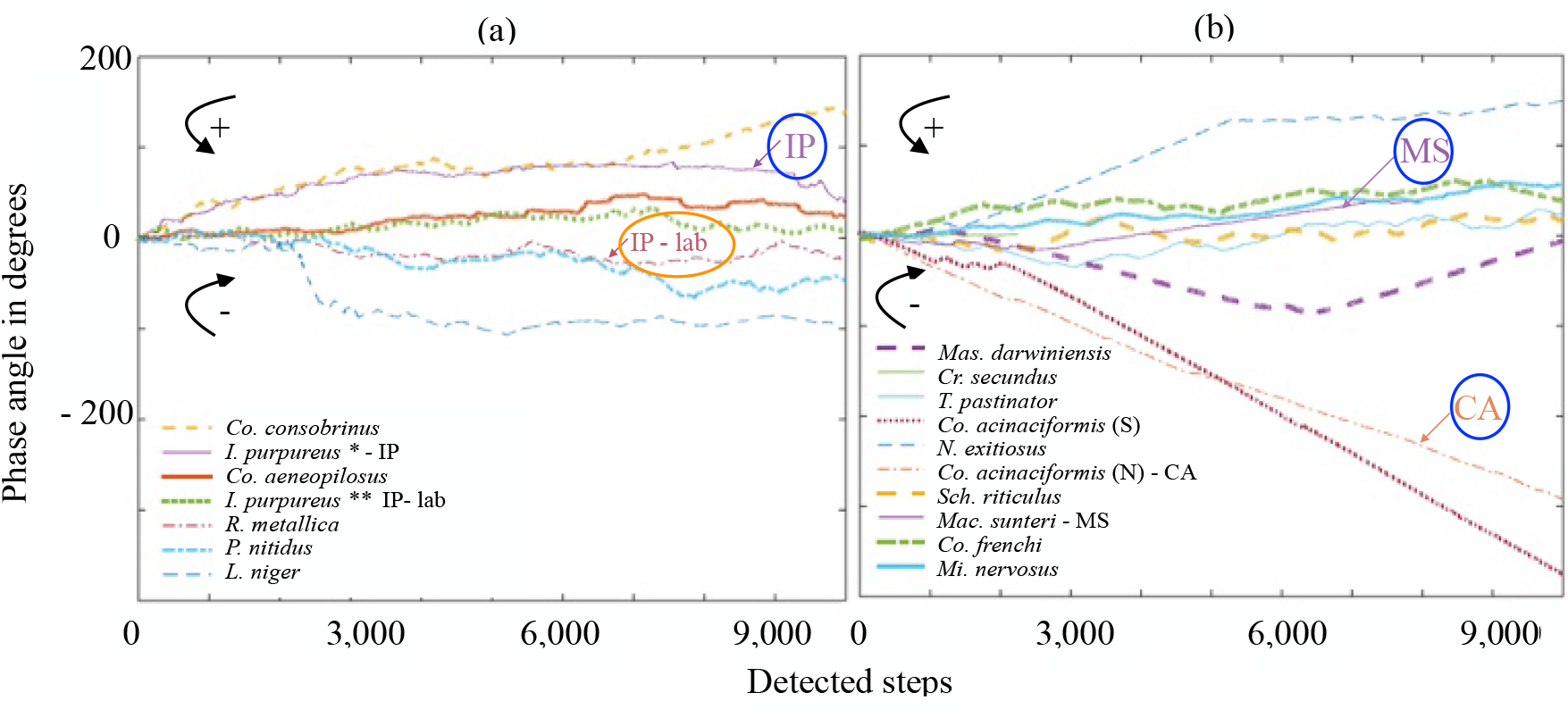
Directional change. Transforming the ants’ and the termites’ walking pattern from Cartesian into polar coordinates and monitoring the phase (in degrees) of the insect walking using the position vector indicates that termites do not change their direction as often and have the tendency to walk continuously in circles, ants’ walking pattern seems to be more erratic (cf. Figure 1). Only the ant *R. metallica* walks more in circles; arrows refer to the direction of motion. The focus species *Iridomyrmex purpureus* (IP), *Coptotermes acinaciformis* (CA), and *Macrognathotermes sunteri* (MS) have been circled in blue for better visibility; the orange-circled species shows the in the laboratory-reared ant *Ir. purpureus* (IP-lab).

Figure 7 depicts motion quantities of different insect species, including the median speed (body-length (bl) per s), median absolute speed (in mm/s), and maximum speed (bl normalised), and the average activity level in %. In Figure 7(a), the median body length in ants is strongly negatively correlated (*ρ* = − 0.92) to the median speed with the smallest ant (3 mm) *L. niger* being the fastest (5.2 bl /s) and the largest (11.9 mm) ant *Ca. consobrinus*, being the slowest (1.8 bl/s). In termites, this correlation is low (*ρ* = − 0.21). Overall, termites are moving slower than ants, relative to their bl, and speeds cluster mostly between 1.8 bl/s to 3.6 bl/s (outlier *N. exitiosus* with 5.0 bl/s) while most ants can be found between 3.3 bl/s and 5.2 bl/s (outlier *Ca. consobrinus* with 1.8 bl/s). The median absolute speed against the median maximum speed (bl/s) for termites shows a positive correlation of *ρ* = 0.67 (ants *ρ* = 0.26), which could potentially be related to the observation made through the video analysis (Figure 7) that termites kept walking at a rather constant speed without standing still.

**FIG. 7.**
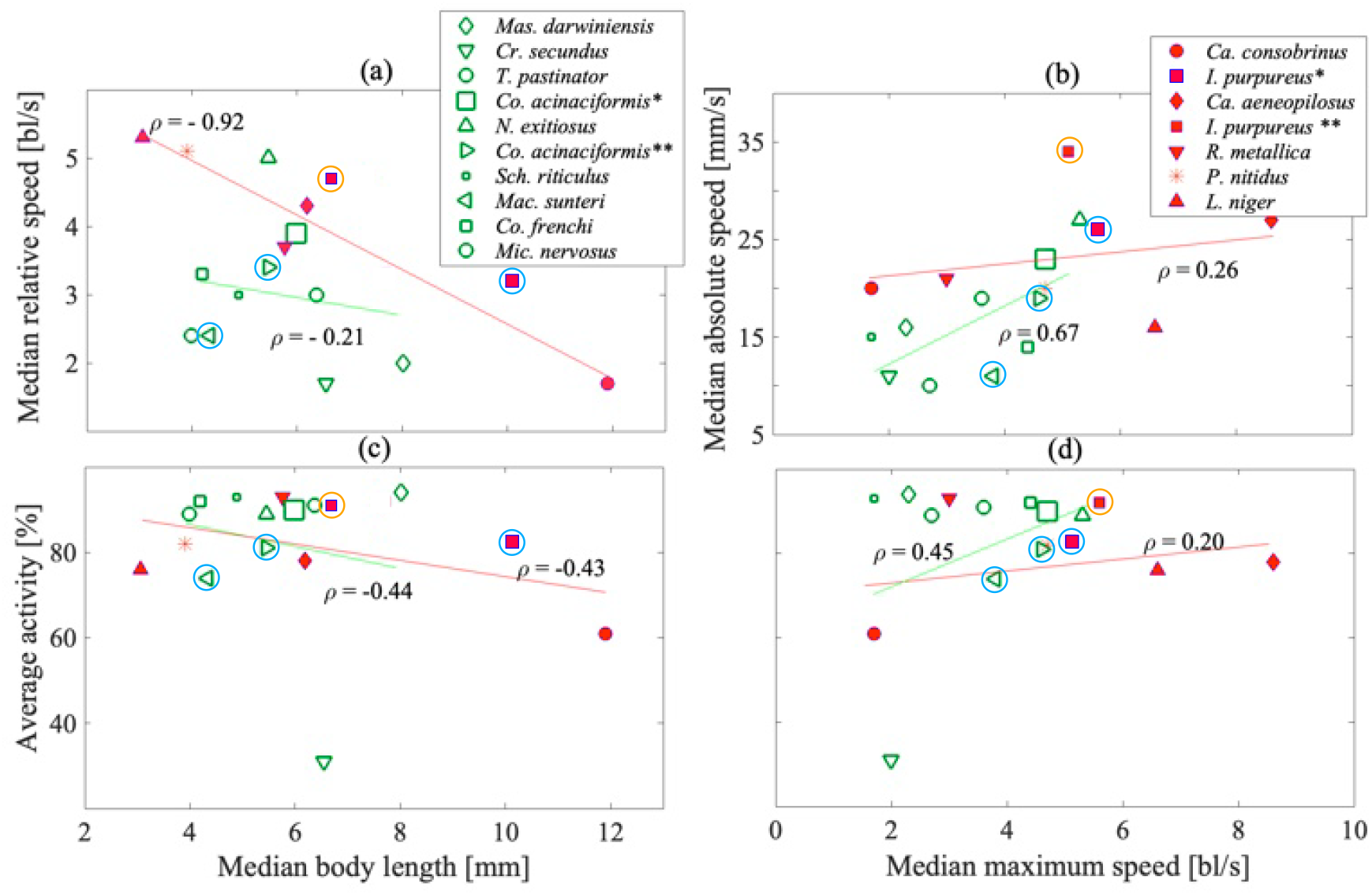
Statistics of ant and termite motion. Extracted post-processed results of the tracking algorithm monitoring ants and termites: (a) the median relative speed (in body length (bl) per second) vs the median body length (mm); (b) the median absolute speed (in mm/s) over the average maximum relative speed (bl/s); (c) the average activity level (in %) over the mean body length and the average activity level over the mean maximum speed. The linear trend for ants and termites is also depicted, as well as the correlation coefficient calculated over all values for ants and termites. For better visibility, blue-circled species indicate the focus species; the orange-circled species shows the in the laboratory-reared ant *Ir. purpureus*.

The quantification over the parameters extracted from the tracking algorithm, that is, the median speed in bl/s and mm/s, the maximum speed, and the average activity, may indicate a connection to the often observed dominant circular motion in termites, see Figure 5. The average activity level (Figure 7 (c)) is similar between ants and termites and lies between 76% for the termite *Mac. sunteri* and 87% for the termite *M. darwiniensis*. The lowest activity of 61% for the ant *Ca. consobrinus*) is considered an outlier.

### B. Vibration analysis

Figure 8 shows the distribution of vibration magnitudes and decay times for different species of ants in (a) and (b) and of termites in (c) and (d). The magnitudes show a strongly positive skewed Gaussian distribution, while the decay times are exponentially distributed. The most platykurtic distribution magnitudes are provided by the laboratory form of *I. purpureus* with values ranging from 6E-5 m/s to 7E-2 m/s. This upper bound is also reached by *C. consobrinus, I. purpureus* (wild form), *C. aeneopilosus*, with relatively high relative frequencies in this area.

**FIG. 8.**
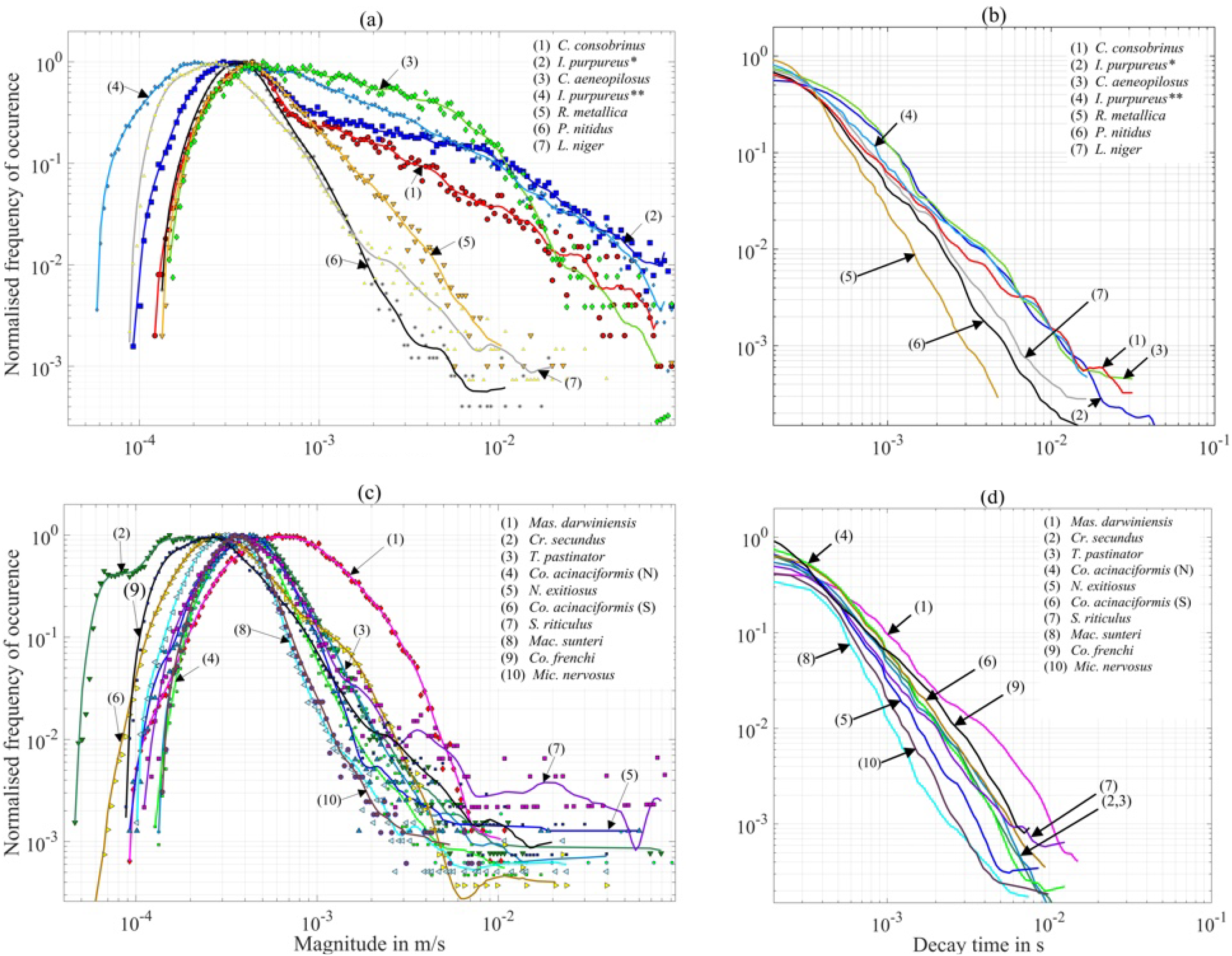
Statistical distributions of enveloper. Median frequencies (markers) and estimated continuous probability distributions over the veneer disc extracted after non-linear filtering using the enveloper for median velocity amplitudes and decay times: (a) and (b) ants; (c) and (d) termites. See Supplementary Materials Figure S2 on how to form the distributions.

Table 1 provides details of the statistics depicted in Figure 8, which were extracted from repeatedly applying the enveloper function to the vibration signals of different ant and termite species.

**TABLE 1.** Ants and Termites - statistics from enveloper function. Surface velocity magnitudes (MAG) and attenuation times (*τ*) as extracted via enveloper function for different ant species (min, median, average, and max values). While the median magnitudes are similar for ants and termites, termites have lower impacts, and their attenuation time *τ* (min and max) is significantly smaller. *I. purpureus\** represents the wild form, which is larger, and *I. purpureus\*\** is the smaller laboratory version of the same species; *Co. acinaciformis\** is the Southern form, which is larger, while *Co. acinaciformis\*\** is the smaller Northern form, collected from Darwin.

| Ant species | length<br>in mm | weight<br>in mg | min(MAG)<br>in $\mu\text{m/s}$ | median(MAG)<br>in $\mu\text{m/s}$ | max(MAG)<br>in mm/s | $\tau_{\text{mean}}$<br>in ms | $\tau_{\text{max}}$<br>in ms |
| --- | --- | --- | --- | --- | --- | --- | --- |
| *C. consobrinus* | 13.5 | 48.3 | 121 | 405 | 16.8 | 31.4 | 204 |
| *I. purpureus* * | 10.0 | 8.6 | 92 | 337 | 45.5 | 41.5 | 178 |
| *C. aeneopilosus* | 8.0 | 11.1 | 146 | 488 | 18.0 | 31.4 | 155 |
| *I. purpureus* ** | 6.5 | 6.7 | 146 | 212 | 39.6 | 16.6 | 77 |
| *R. metallica* | 5.0 | 5.4 | 58 | 405 | 32.5 | 4.7 | 47 |
| *P. nitidus* | 3.0 | 1.4 | 134 | 353 | 15.5 | 11.4 | 107 |
| *L. niger* | 2.5 | 1.0 | 88 | 267 | 16.2 | 16.4 | 123 |
| Termite species | length<br>in mm | weight<br>in mg | min(MAG)<br>in $\mu\text{m/s}$ | median(MAG)<br>in $\mu\text{m/s}$ | max(MAG)<br>in mm/s | $\tau_{\text{mean}}$<br>in ms | $\tau_{\text{max}}$<br>in ms |
| *Mas. darwiniensis* | 10.75 | 26.9 | 171 | 644 | 4.3 | 1.91 | 14.99 |
| *Cr. secundus* | 6.75 | 8.0 | 153 | 405 | 1.8 | 1.11 | 10.35 |
| *T. pastinator* | 5.5 | 5.9 | 133 | 425 | 1.9 | 1.10 | 9.44 |
| *Co. acinaciformis* * | 5.3 | 3.9 | 127 | 387 | 1.3 | 0.97 | 12.49 |
| *N. exitiosus* | 4.5 | 5.0 | 106 | 426 | 1.4 | 0.73 | 8.6 |
| *Co. acinaciformis* ** | 4.35 | 4.8 | 111 | 280 | 2.2 | 1.06 | 9.44 |
| *S. riticulus* | 4.2 | 1.9 | 111 | 387 | 2.1 | 1.06 | 12.46 |
| *Mac. sunteri* | 4.2 | 1.9 | 140 | 370 | 1.1 | 0.59 | 7.49 |
| *Co. frenchi* | 4.1 | 1.7 | 88 | 255 | 1.6 | 0.60 | 7.54 |
| *Mic. nervosus* | 3.85 | 2.3 | 101 | 322 | 1.0 | 0.74 | 9.89 |

As expected, the average minimum, median, and maximum amplitudes depend as expected on the insect’s size, hence their weight (see Figure SI for graphical data). Even though the average median magnitudes are similar in the order of magnitude as determined by Oberst et al. (2017) [15], significant differences to absolute maximum values within bins of the distribution belonging to the right tail can be observed. The average minimum magnitudes for *Rhytidoponera metallica* are the smallest among the ants studied, and also the mean decay time is the shortest. This species is more basal than the other species studied, and mostly hunts alone; findings from [15] also indicate that this species is quieter than other ant species. The average mean attenuation times are found to be of the same order of magnitude as observed by Oberst et al. (2015) [19], while large differences in average maximum attenuation times can be observed. *R. metallica* has the lowest value in their mean decay time 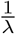. Similar to ants, the average magnitudes in termites decrease with size (weight) but are less broad and less positively skewed (Fig. 8); and, in accordance with [15], much lower values (a factor of 100) than those found in ants. All distributions look similar except for *M. darwiniensis*, which is shifted to the right while *Cr. secundus* is shifted to the left. The fastest vibration attenuation time is found in *Mac. sunteri* followed by *Co. frenchi*. Figure S3 in the Supplementary Materials summarises and correlates the results of the enveloper function statistics into two graphs and connects them to the weight of the insects (Figure SI 1). Using a polynomial fit in Matlab, the data points of median velocity amplitude vs median decay time fit a near quadratic function of median vibration substrate response amplitude in ants (*v* = 4.5*E* − 3*t*^1.86^ mm/s) and form a square-root-like behaviour for termites (*v* = 7*E* − 4*t*^0.49^ mm/s), *t* being the median decay time extracted from Fig 8.

#### 1. Power spectral density estimations

Figure 9 shows the median peak power spectral density of various vibration velocity measurements to identify the frequency range of resonances involved in the response behaviour of the veneer disc. For both ants and termites, the highest resonance scatters between 0.7 and 1.5 kHz; ca. 0.2 mm/s is the reference background noise level, indicating that termite footstep response signals are often close to or buried in noise. Termites excite many resonances, and usually, the peak PSD estimate for termites is lower than the background noise level except for *Co. acinaciformis*. Ants are also two orders of magnitude louder than termites. Figure S4 in the Supplementary Materials shows the integrated power spectral density for all species studied, which confirms findings of [15] and clusters ants and termites into two groups.

**FIG. 9.**
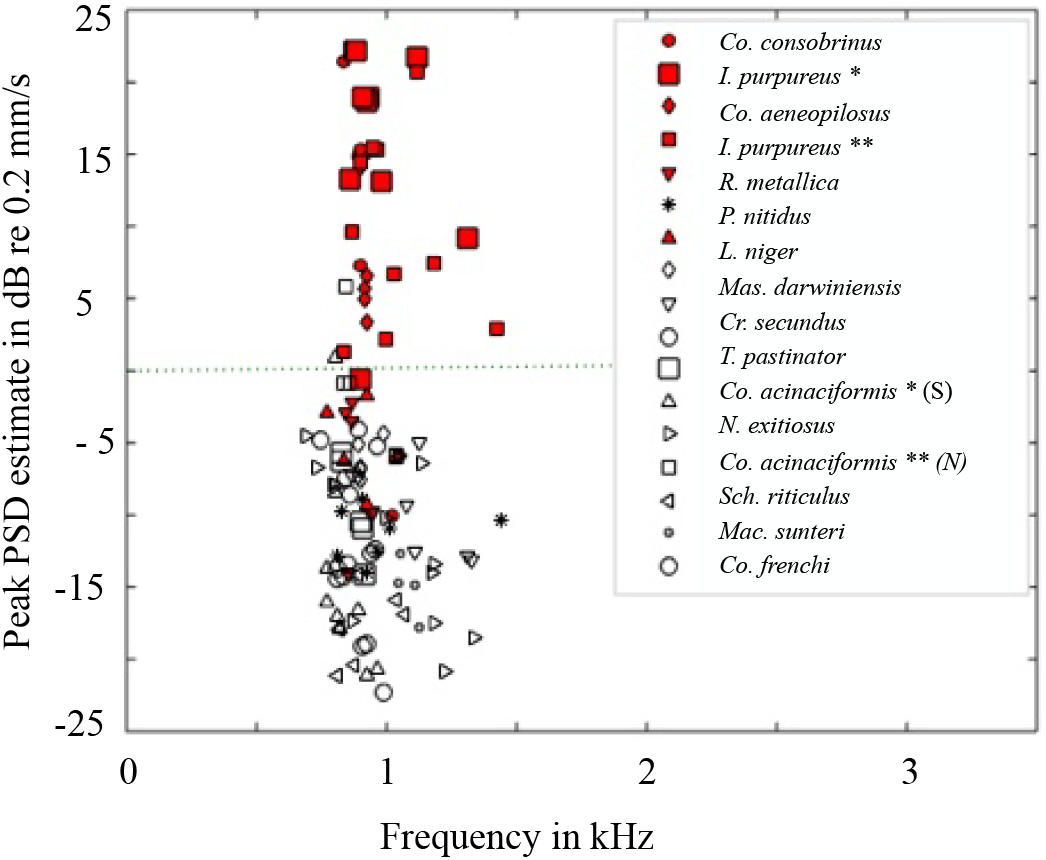
Frequency domain analysis, vibration signal. Peak power spectral densities over frequency. Ants’ peak PSD estimates are up to two orders of magnitude higher than those of termites.

### C. Highly Comparative Time Series Analysis

Applying HCTSA to cases A-C (see Supplementary Materials, Figures S5 to S7) shows that HCTSA can distinguish between different analytical benchmark systems. These systems include the background noise signal and white noise, filtered and unfiltered data, termites and ants (*Co. acinaciformis* and *Ir. purpureus*) response signal data. All of these systems show that the HCTSA analysis works well, cluster quality is high, and the probability that clustering has occurred by chance is very low.

Figures 10, 11, and 12 show the application of HCTSA to cases D, E, and F.

**FIG. 10.**
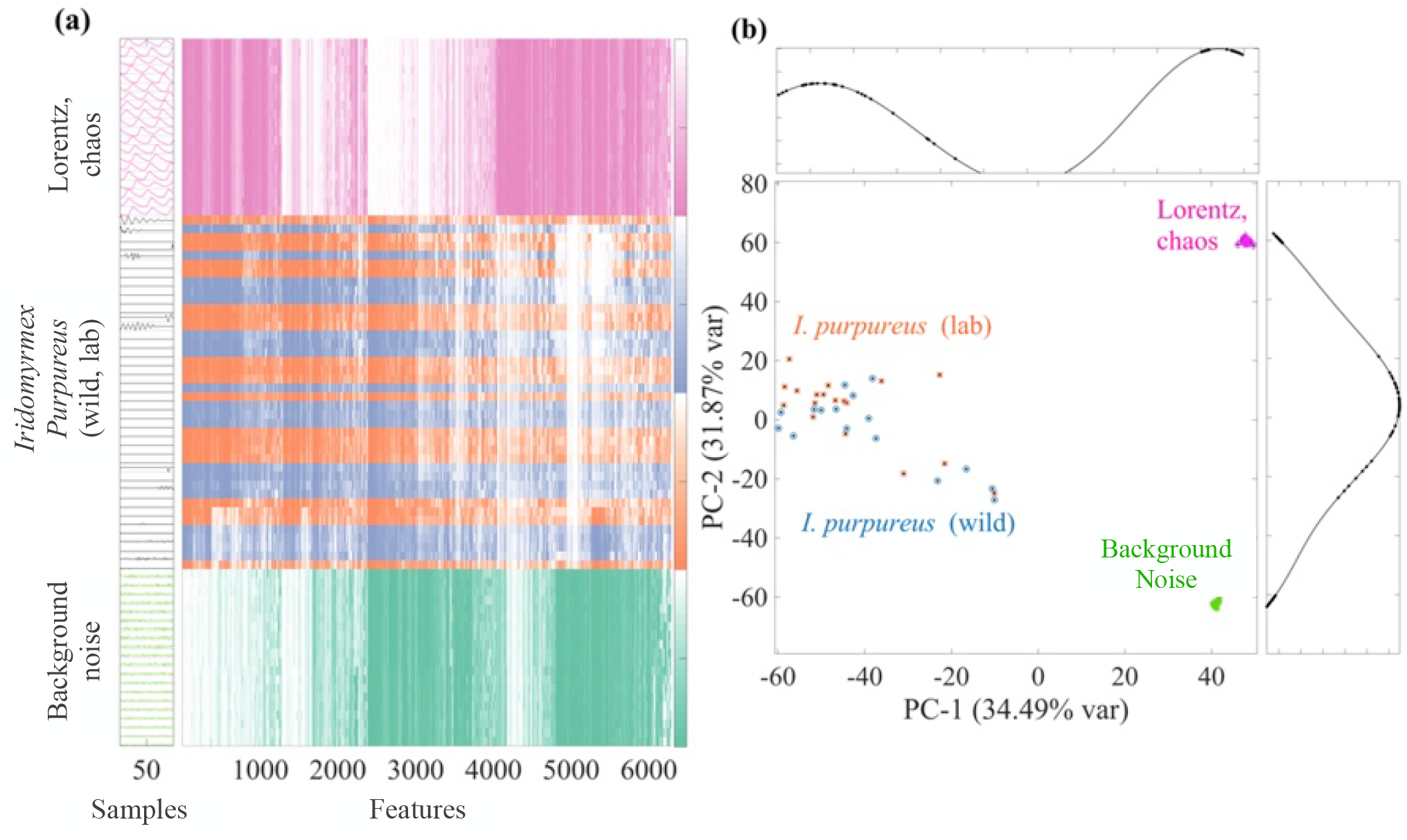
Case D: Size effect. a) shows the HCTSA of the Lorentz system, laboratory and wild form of *I. purpureus*, and the background noise. b) Principal component analysis of the signals analysed; PC-1 captures the maximum variance in the data. PC-2 captures the next highest variance, subject to being orthogonal (uncorrelated) to PC-1. The results show that the main features of the wild form can not be distinguished from those of the laboratory colonies. Scale on the y-axis in b) on the inserts is normalised to one.

**FIG. 11.**
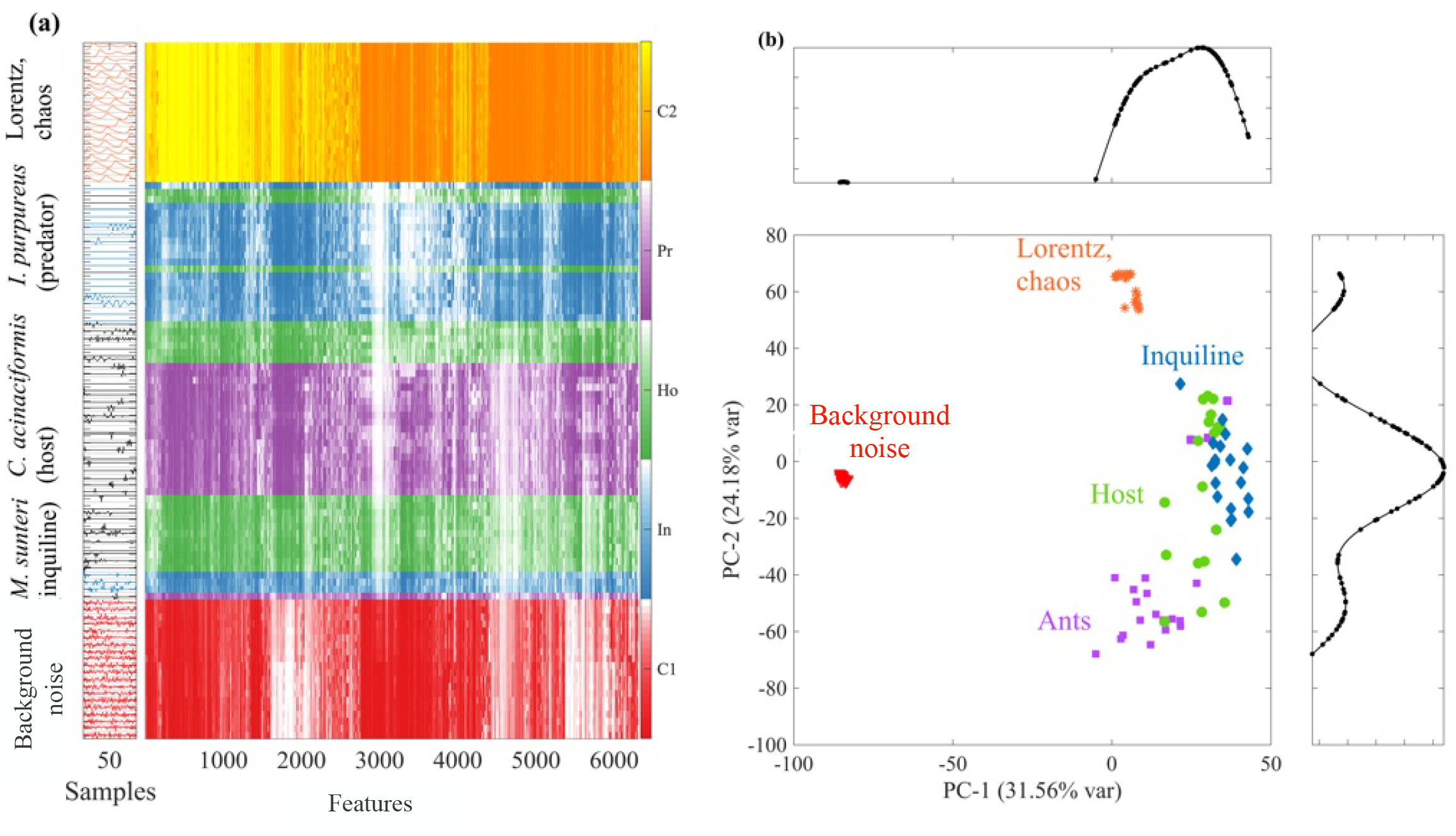
Case E: Predator-prey/host-inquiline. a) shows the HCTSA of the Lorentz system, wild form of *I. purpureus* as an ant predator, *C. acinaciformis* as termite prey/host, *M. sunteri* as a termite prey/inquiline (to host) and the background noise. b) Principal component analysis of the signals analysed; PC-1 captures the maximum variance in the data. PC-2 captures the next highest variance, subject to being orthogonal (uncorrelated) to PC1. Termites and ant signals (laboratory form of *Iridomyrmex purpureus* are separable in their features, but the inquiline’s main features spread within its host’s signal. Scale on y-axis in b) on inserts is normalised to one.

**FIG. 12.**
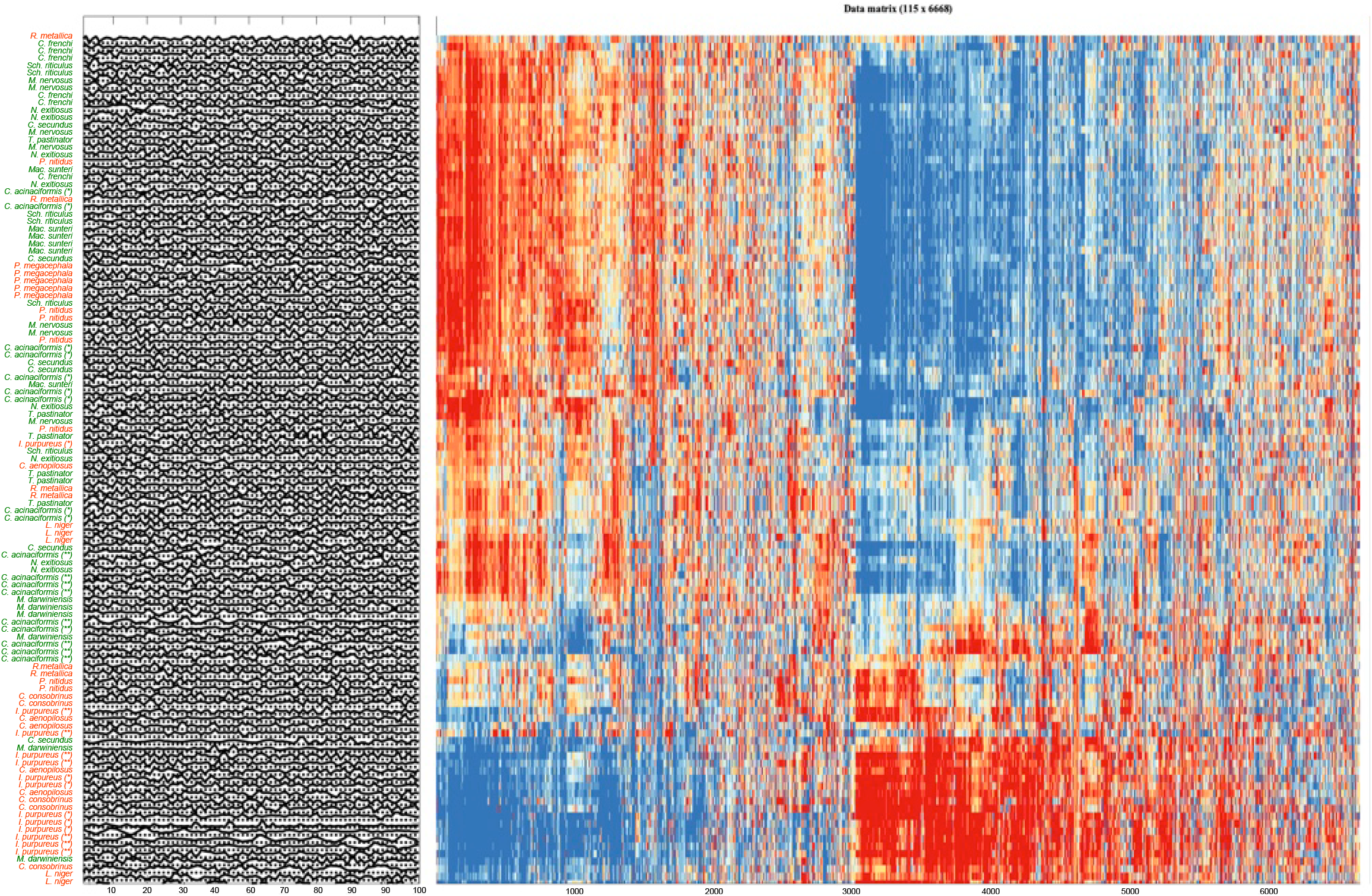
Case F: All species comparison. On the left, various samples of ants (red) and termite (green) species are listed. In the HCTSA matrix a red colour indicates a *high correlation*, and a blue colour indicates *low correlation*. Signals of termite species relate more to the first 1,000 features, whereas ants have a stronger correlation for features showing up after 4,500 to about 5,500.

Figure 10 compares the laboratory and the wild form of *Ir. purpureus*. One can see that the features of signals of the ants studied seem to be independent of the individual’s size or weight (cf. Figure S1); the laboratory and the wild form have, after normalisation with the maximum amplitude, identical feature clusters. The signals are also well separated from the Lorenz chaotic data and the background noise. Size dependence, therefore, does not play a major role in the HCTSA features, especially since termites and ants can be well separated for amplitude-normalised filtered data (Figure S6). The quality of the clustering for Lorenz, lab, and wild form of *I. purpureus* and noise is *Q*(*c*) = 100%, 11%, 15% and 100% respectively.

The exact probability relative to the probability of clustering by chance is effectively zero. Even using precise factorial calculations, the probability is so small that it rounds to zero in floating-point arithmetic. This confirms that the event is virtually impossible to occur by random chance. Conducting a Chi-Square goodness-of-fit test in Matlab gives a *p*-value below the floating-point accuracy, which is effectively zero; hence, the clustering as presented here is unlikely to be a result of chance.

In Figure 11, the three species *Ir. purpureus* (predator), *Co. acinaciformis* (prey/host) and *Mac. sunteri* (inquiline/prey) are studied. Again, similar to Figure S7, ants and termites can be separated using the PCA. However, only using the ranks in the HCTSA matrix, the inquiline and the host can be clustered as shown through large connected areas of blue and green. The clustering quality for Case E is then is *Q*(*c*) = 100% (Lorentz), 35% (Predator), 58% (Host), 90% (Inquiline), and 100% (Background noise), respectively. Some mixing occurs, which might be due to the high similarity of some signals. More features than the two principal components are required to distinguish between the host and the inquiline species.

The exact probability *P* (*k, m, n*) is again effectively zero. and the Chi-Square goodness of fit test in Matlab also gives a *p*-value below the floating point accuracy, which is effectively zero.

Figure **??** illustrates that termite and ant signals can be clustered based on their feature profiles. While some uncertainty remains, termite signals predominantly correlate with features in the range up to ID 1,000 (strong correlation shown in red), whereas ant signals show strong correlation with features between ID 4,500 and 5,500.

The feature range from 1 to 1,000 is dominated by mutual information (AMI) with 289 occurrences, alongside information-theoretic measures (248), correlation-based features (248), stationarity indicators (192), distributional statistics (86), and entropy measures (68) [47].

These features suggest that termite signals are structured, exhibit temporal dependencies, and are statistically stable over time. Their dynamics appear deterministic, possibly reflecting low-dimensional chaotic behavior that is synchronized and predictable, akin to controlled system outputs or biological rhythms.

In contrast, the feature range from 4,500 to 5,500 is rich in nonlinear dynamics (346), pre-processing-sensitive techniques (502), Gaussian process models (102), and prediction-oriented methods [47].

These features imply that ant signals are higher-dimensional, nonstationary, and more complex, often influenced by noise, trends, or artifacts. Their statistical properties evolve, and the underlying dynamics are less synchronized and more variable, requiring more sophisticated modeling approaches.

The results in terms of clustering quality and probabilites are provided below in Table II.

**TABLE II.** Comparison of theoretical probabilities for ants and termites. Overview of results according to clustering of ants and termites. While the clustering score is low, the probability that the accumulation of samples in Figure 12 is based on chance is unlikely.

| Ants | % $Q(c)$<br>see Eq. 3 | $P(k, n, m)$<br>see Eq. 8 | Termites | % $Q(c)$<br>see Eq. 3 | $P(k, n, m)$<br>see Eq. 8 |
| --- | --- | --- | --- | --- | --- |
| *R. metallica* | 6% | 1.39000000% | *C. frenchi* | 28% | 0.00220000% |
| *I. purpureus* * | 28% | 0.01620000% | *Sch. riticulus* | 6% | 0.00075000% |
| *I. purpureus* ** | 57% | 0.00800000% | *M. nervosus* | 4% | 0.00260000% |
| *P. nitidus* | 2% | 0.00730000% | *N. exitiosus* | 3% | 0.00780000% |
| *P. megacephala* | 100% | 0.00007240% | *C. secundus* | 3% | 0.01370000% |
| *L. niger* | 40% | 0.00081000% | *T. pastinator* | 3% | 0.01370000% |
| *C. aenopilosus* | 4% | 0.00170000% | *C. acinaciformis* * | 4% | 0.00001200% |
| *C. consobrinus* | 8% | 0.00001640% | *C. acinaciformis* ** | 25% | 0.00000185% |
|  |  |  | *Mac. sunteri* | 44% | 0.00021000% |
|  |  |  | *M. darwiniensis* | 25% | 0.00110000% |

Although some clustering quality values, *Q*(*c*), appear quite low, in every case the probability *P* (*k, n, m*) is significantly smaller than *Q*(*c*). This indicates that the observed results are unlikely to be due to random chance, as a stochastic model based on neighboring elements provides a realistic measure of randomness.

## IV. DISCUSSION

Termites and ants have been engaged in a predator-prey arms race spanning millions of years [15, 21]. Oberst et al. [15] observed that termites (social cockroaches) move significantly more quietly than ants, and their alarm signals appear to mimic the footsteps of their predators. Of particular interest is the relationship between the predatory meat ant *Ir. purpureus* and its termite prey *Co. acinaciformis*, which itself shares a host-inquiline relationship with *Mac. sunteri*. The intricate dynamics of this predator-prey and host-inquiline system, as depicted in Figure 1 could be based on vibrations of substrates induced by walking insects. Hence, this study was focussed on the characteristics of the walking vibrations of six ant and ten termite species and on exploring whether these vibration signals shed some light on the predator-prey and host-inquiline relationships.

Framed in the context of the long-standing predator–prey arms race [15] between ants and termites, these results provide insight into how locomotion-derived substrate vibrations may function as ecologically relevant signals rather than incidental by-products of movement. While ants, termites, and inquilines share broadly similar locomotory patterns, our findings show that the resulting vibrational signatures differ systematically in intensity, temporal structure, and predictability, in ways that more align with their ecological roles. Termites, including host and inquiline species, tend to move more slowly and quietly, yet generate more structured, rather harmonic (low dimensional) vibration patterns, whereas ants produce louder, more irregular (high-dimensional), and less predictable and rather random-looking signals. These differences are consistent with our hypotheses of reduced detectability and stealth in prey and cohabiting species [14], and enhanced signalling or unavoidable vibrational conspicuousness in predators, without requiring explicit assumptions of adaptive mimicry.

Vibration responses produced by walking or running on a substrate, which can be eavesdropped on by predators, competitors, or inquilines, arise from a complex interplay of various mechanisms. These include the kinematics and general physiology of the legs, whole-body motion, and the interaction of the substrate with the abdomen. Additionally, the insect’s behaviour, which is mode-dependent and situational, as well as the adaptation of the feet to the substrate, plays crucial roles. The feet contribute through friction and adhesion, which influence the gait pattern [58].

Past studies have compared the kinematics of ants and cockroaches (*Blaberus discoindalis*), which exhibit different running speeds. For example, *Blaberus discoindalis* can run up to 24 body lengths per second (bl/s) [23], while *Formica pratensis* and *Formica polyctena* have been measured at 9 bl/s and 12 bl/s, respectively [24]. Although both ants and cockroaches show rhythmic gait fluctuations, differences in movement control were observed, likely due to ants inhabiting more rugged environments, whereas cockroaches are adapted to flat terrain [24].

Our analysis, however, reveals that in the confined area of the arena used, the maximum median speeds of ants (5.2 bl/s) and termites (5.0 bl/s) were much lower than those reported previously [23, 24]. While we used different species, the experimental setup may have influenced the observed patterns and behaviours. The laboratory form of the meat ant, which was reared in the lab (approximately 65% of the size of the wild form), was relatively faster and only slightly more active than the wild form. Conversely, the largest ant (*Ca. consobrinus*) was the slowest and least active. Oberst et al. [15] linked changes in behaviour to *nesting behaviour*, related to the density of individuals in an area, leading to higher rates of quasi-static behaviour, with increased trophallaxis and grooming among nest mates. A similar effect may have been observed here, where the overall space available appeared to determine activity levels and speed.

Regarding vibration response levels, Oberst et al. [15] showed that ants produce higher-intensity vibrations than termites during motion. We found that almost all termites were quieter than ants, except for *Co. acinaciformis*, which are louder than the background noise levels (Figure 9). *Co. acinaciformis* produced vibrations slightly above the background noise level, while ant species like *R. metallica* and *L. niger* were generally quieter. Spectral analysis revealed that termites excite more resonances within the vibration frequency range of 740 to 1,500 Hz, which contains the first and second vibration modes of the veneer disc [17]. While ants mostly exhibited erratic, mixed walking behaviour, dominant termite species, which are abundant and live in large colonies, moved in a more circular pattern [57]. *Mic. nervosus* and *Mac. sunteri* moved in irregular, seemingly random patterns, covering large areas of the arena, while their host *Co. acinaciformis*^∗ ∗^ (S) displayed a more circular movement pattern. Among ants, only *R. metallica* consistently moved along the edge of the arena, yet it remained one of the quietest species, despite being larger compared to *P. nitidus* and *L. niger* (Table 1), confirming the findings of [15].

The median speeds of termites in both bl/s and mm/s were observed to be lower than those of ants. Additionally, termite speed showed a much weaker negative correlation with body length (≈ *ρ* = − 0.21) compared to ants (≈ *ρ* = − 0.92). Activity levels and the correlation between activity levels and body length were similar for both ants and termites (≈ *ρ* = − 0.43). Although termites are social cockroaches, unlike *Blaberus discoidalis*, they typically move within self-built clay corridors, either in their nests or at foraging sites [3, 59]. While providing ideal environmental conditions (humidity, temperature), likely these structures have also ideal topological designs i.e. curvatures, slopes, and smooth inner surfaces, with optimised pathways for energy efficient and secure movements [3, 59, 60].

The fact that termites move slower and quieter yet produce more harmonic vibrations during movement in the arena than ants may be related to their unique movement strategies and how they leverage their physiology for specific behaviours. Reinhardt et al. (2009) [24] observed that the lateral component of the ground reaction forces in *Formica* ants exhibited small oscillations at higher frequencies, and the vertical force component also oscillated, which differed from observations in cockroaches. These oscillations, along with the role of the abdomen, are not captured in current analytical models (SLIP/BSLIP, LLS). It is suggested that the ant’s gaster plays a similar role in the fore-aft direction as gravity does during climbing. In contrast, Full et al. (1991) [23] reported that ground reaction forces in the cockroach *B. discoidalis* were unimodal, with an even vertical force distribution across all legs, and the time evolution of frontal and lateral components was simpler, without significant braking forces generated by the abdomen.

Ants have longer legs relative to their body size, while the gaster in termites is generally closer to the ground. Unlike cockroaches, termites are eusocial insects that use trail-following pheromones to communicate chemically with nestmates. These pheromones are deposited along the path during movement, secreted through the sternal glands located in the posterior metasoma, or gaster [61]. Due to this additional contact, the vibrations of the veneer disc may become further attenuated, similar to how a drum’s sound is muted by the membrane. The contact (or breaking force in ants) may be more pronounced in termites, especially relative to their speed, which could lead to a shorter attenuation time and consequently a lower overall vibration level. Since cockroaches have less abdominal surface-body contact, the absence of this braking force may naturally result in faster movements [23, 24].

The excitation of the first vibration mode, which has a single nodal circle, generates a higher absolute vibration amplitude than the second mode, which has an additional nodal line running across the diameter [62]. However, the specific modes responsible for the higher vibration amplitude may be less significant if termites adjust their behaviour to minimize vibrations, moving across an area in the quietest possible manner.

Movement in a mixed pattern appears more random, while circular motion around the edge of the arena likely excites primarily the first vibration mode, with a single circumferential nodal line. In contrast, the mostly random walking pattern seems to excite additional modes of the setup, distributing the energy more widely. This distribution may lead to a quieter response and a reduced ability to localize vibrations. Both random and synchronized circular movements are well understood in terms of collective behaviour in animals, including arthropods [5, 63]. Whether similar behaviour can be observed in groups of ants and termites remains an open question.

We demonstrate that the highly comparative time series analysis (HCTSA) can successfully distinguish (cluster) different benchmark systems (Case A): a sinusoidal signal, a Lorenz system in both a nonlinear periodic regime and a chaotic regime, as well as recorded background noise and Matlab-generated white noise (Figure S5). Additionally, HCTSA effectively clusters background noise from the original *Co. acinaciformis* recordings and their *ghkss*-filtered time series (Case B, Figure S6). When providing HCTSA-filtered time series for *Ir. purpureus, Co. acinaciformis*, Lorenz chaos, and background noise recording, all groups are separated, with some overlap between ants and termites (Case C, Figure S7).

We further show that the vibration velocity signals contain features distinct from those dominated by size and weight, as represented by the data in Table 1. The laboratory and wild forms of *Ir. purpureus* mix well in their feature sets, and comparison of the HCTSA matrix (Case D, Figure 10) indicates that, after amplitude normalization, other features of the vibration response are strikingly similar, if not identical, for the species. Comparing the clusters between the inquiline *Mac. sunteri*, its host *Co. acinaciformis*, and its predator *Ir. purpureus* shows a clear separation (Case E, Figure 11), with this distinction being especially evident in the feature matrix. The separation is much clearer than the overlap observed between the laboratory and wild forms of *Ir. purpureus* (Figure 10).

Computing all species using the HCTSA, it is not possible to use just two principal components, and more features are required. Also, clustering becomes more diverse and has outliers. Overall, similarity in walking between ants and termites, as well as different behaviour during motion, as well as the vibration caused at different locations, might lead to inaccuracies in labelling the data, and a finer breakdown of the behaviour would be required. This, however, would need a close-up video recording to see more detail. Overall, the results show that more features are required for clustering.

When studying the features and their higher classification, features mainly found in termites suggest higher structures pattern, with temporal dependencies, and statistically stable signals over time. Their dynamics appear deterministic, possibly reflecting low-dimensional chaotic behavior that is synchronized and predictable, akin to controlled system outputs or biological rhythms [47].

The measures which were found to be dominant in ant species seem to be nonstationary, and more complex, often influenced by noise, trends, and with some sorts of artifacts. Their statistical properties evolve, and the underlying dynamics are less synchronized and more variable, requiring more sophisticated modeling approaches; this indicates higher randomness and higher dimensionality of the signals [47]. However, what pattern exactly exist and how to model their walking response pattern in an ideal way e.g. using bio-inspired insect gait as excitation function in the noise-control engineering principle [2, 64, 65], remains an open question which is to be explored in the future.

From the HCTSA vibration signal analysis, it appears that *Mac. sunteri* shares similarities with the signals of *Co. acinaciformis*, with some features closely resembling those of its host. This similarity could be attributed to both species being termites, which results in their signals being distinct from those of *Ir. purpureus*. However, the fact that they still cluster separately suggests that the inquiline emits vibration signals through the substrate that share similar yet distinct features compared to those produced by the host during walking. The reduction to two principal components further indicates that the inquiline signal, and more generally the termite walking signals, align more closely with deterministic chaotic features of the Lorenz system. This could be coincidental, but it may also indicate complex walking behaviours that mimic randomness to appear less predictable, likely driven by varying motion in the arena and different modes being excited over time. However, whether the insects synchronise their walking, and if this walking is in indeed deterministic and chaotic and not following a Lévy flight [28] will need to be confirmed in future studies.

Nevertheless, further work is required to determine how foraging decisions under threat are shaped by the vibration-transmission properties of the substrate. Because the observed clustering and comparison with a stochastic model are primarily correlative, their interpretation should be treated cautiously. Collective behaviour may also influence how ants and termites synchronise or modulate their movement patterns. It is therefore important to validate these findings—particularly the interactions between *Coptotermes acinaciformis* and *Macrognathotermes sunteri*, through targeted bioassays to strengthen ecological and evolutionary conclusions and to establish causation.

Such experiments would clarify the dynamic processes underpinning cohabitation, defence, and foraging, which may themselves depend on physiological constraints and species-specific sensory limits. This requires dedicated bioassays and expanded arena studies, similar in design to those presented here but involving larger numbers of individuals. Ultimately, field studies will also be essential, as walking behaviour changes in captivity and with spatial restriction (cf. [15]). Incorporating behavioural feedback loops [64] will further reveal how species such as *M. sunteri* adjust their walking patterns in response to increasing proximity or activity of *C. acinaciformis*.

The development of a more sensitive force plate to study the gait patterns of termites in comparison to ants is underway, but this remains an outstanding research task [66–68]. Here also the proper characterisation of the bioassay from its structural aspect, considering different vibration modes and increasing the spatial sampling, is an important, yet outstanding task. The precise effect of abdominal body contact during walking, and the differences between ants and termites, should be further explored using a synchronized high-speed camera, linking vibration and force pattern with kinematics. This approach would also enable the study of insect joint trajectory patterns, supporting the development of quiet, insect-inspired autonomous robotic walkers for use in disaster response and space exploration.

## DATA AVAILABILITY

All data analysis methods are available in the manuscript. Any questions should be addressed to the corresponding author.

## AUTHOR CONTRIBUTIONS

A CRediT authorship contribution statement is provided in the following. **SO**: Conceptualisation, Methodology, Investigation, Formal analysis, Data curation, Programming/Scripting and Software, Visualization, Validation, Writing – original draft, Resources, Funding acquisition, Project administration, Supervision. **JL**: Methodology, Investigation, Writing - review & editing, Resources, Funding acquisition. **TAE**: Investigation, Resources, Funding acquisition

## ACKNOWLEDGEMENTS

The authors acknowledge a license for scientific activities under the Nature Conservation Act 2014 within the ACT Tidbinbilla Nature Reserve (S.O. permit TS20188).

